# Fat2 polarizes Lar and Sema5c to coordinate the motility of collectively migrating epithelial cells

**DOI:** 10.1101/2023.02.28.530349

**Authors:** Audrey Miller Williams, Sally Horne-Badovinac

## Abstract

Migrating epithelial cells globally align their migration machinery to achieve tissue-level movement. Biochemical signaling across leading-trailing cell-cell interfaces can promote this alignment by partitioning migratory behaviors like protrusion and retraction to opposite sides of the interface. However, how the necessary signaling proteins become organized at this site is poorly understood. The follicular epithelial cells of *Drosophila melanogaster* have two signaling modules at their leading-trailing interfaces—one composed of the atypical cadherin Fat2 and the receptor tyrosine phosphatase Lar, and one composed of Semaphorin 5c and its receptor Plexin A. Here we show that these modules form one interface signaling system with Fat2 at its core. Trailing edge-enriched Fat2 concentrates both Lar and Sema5c at cells’ leading edges, likely by slowing their turnover at this site. Once localized, Lar and Sema5c act in parallel to promote collective migration. Our data suggest a model in which Fat2 couples and polarizes the distributions of multiple effectors that work together to align the migration machinery of neighboring cells.

## Introduction

Epithelial cells migrate collectively during animal development, wound healing, intestinal turnover, and cancer metastasis^1–5^. To do so, they must polarize within the epithelial plane at both the individual and tissue scales. At the individual scale, cells polarize along a leading-trailing axis. Protrusion and adhesion formation are biased to cells’ leading edges, and contractility and adhesion removal to their trailing edges, much as in cells migrating solo^6–8^. At the tissue scale, cells throughout the epithelium are polarized such that their leading edges preferentially point in the direction of migration, and trailing edges in the opposite direction, a form of planar cell polarity. At the intersection of these scales are the cell-cell interfaces that link the trailing edge of one cell to the leading edge of the cell behind. These leading-trailing interfaces can act as sites of biochemical or mechano-chemical signaling that polarize motility behaviors across the interface^9–12^ However, we know little about how the necessary signaling proteins become organized along these interfaces.

The rotational migration of the follicle cells in *Drosophila melanogaster* has proven to be a fruitful system for identifying signaling mechanisms that coordinate epithelial cell movements. Follicle cells are somatic cells of the egg chamber, the multicellular structure within the ovary that gives rise to an egg. They form a continuous monolayer epithelium around a central cluster of germ cells, and they are surrounded in turn by a basement membrane extracellular matrix that encapsulates the entire egg chamber. The follicle cells’ apical surfaces adhere to the germ cells, and their basal surfaces adhere to and crawl along the basement membrane^13^. Migration in this topologically-closed configuration causes the entire egg chamber to rotate within the stationary basement membrane. This motion changes the basement membrane’s structure, ultimately helping give the egg its elongated shape^13–16^. Follicle cell migration requires WAVE complex-dependent lamellipodia, which are polarized to the leading edge of each cell and planar-polarized across the epithelium^14,17^ (Fig. 1A,B). Polarity emerges tissue-autonomously, without input from extrinsic directional cues. This simplifying feature allows us to more easily isolate the contribution of within-group coordination to collective migration. It likely also makes these cells particularly reliant on such coordination for movement.

**Figure 1:**
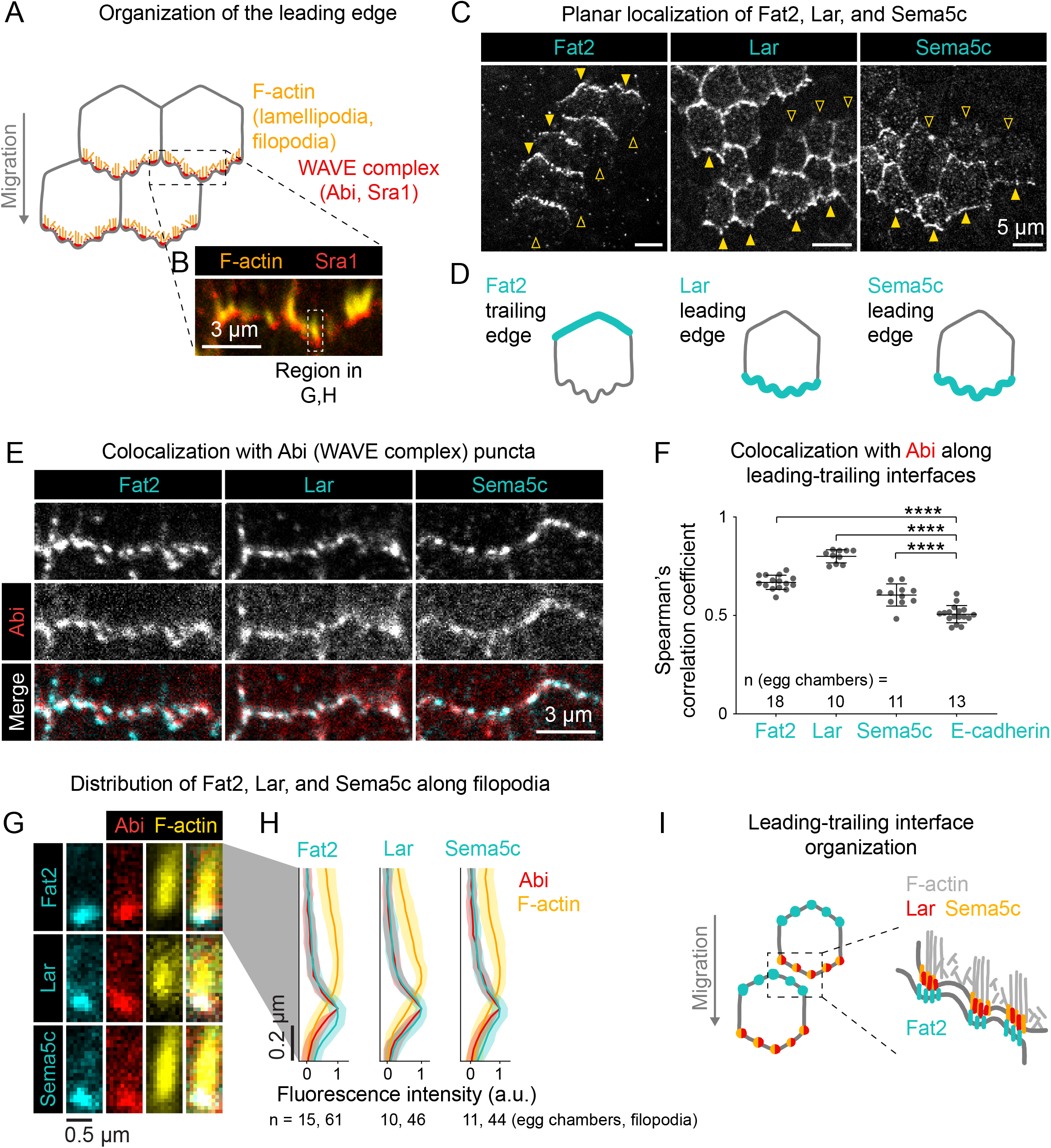
Introduction to the organization of Fat2, Lar, and Sema5c at the follicle cell basal surface. **A**, Diagram of protrusion organization at the basal surface of four follicle cells. The WAVE complex (including subunits Abi and Sra1, labeled in this study) is enriched at the tips of filopodia and lamellipodia along the leading edge of each cell. **B**, Image of the leading edge of one cell with Sra1 (Sra1-GFP) and F-actin (phalloidin) labeled. Filopodia, including a boxed example, are the most prominent F-actin structures. **C**, Images of the basal surfaces of epithelia with mosaic expression of Fat2-3xGFP, Lar-3xGFP, or Sema5c-3xGFP, showing the cell side to which each is polarized. Arrows point to leading-trailing interfaces at the boundary between 3xGFP-labeled and unlabeled cells. Arrows are filled where the labeled protein is enriched and hollow elsewhere. **D**, Diagrams of Fat2, Lar, and Sema5c localization at the basal surface, as shown in (C). Fat2 is polarized to the trailing edge; Lar and Sema5c to the leading edge. **E**, Images of leading-trailing interfaces labeled with Abi-mCherry and Fat2-3xGFP, Lar-3xGFP, or Sema5c-3xGFP. **F**, Plot of colocalization (Spearman’s correlation coefficient) between Abi-mCherry and Fat2-3xGFP, Lar-3xGFP, Sema5c-3xGFP, or E-cadherin-GFP along leading-trailing interfaces. Fat2, Lar, and Sema5c all colocalize with Abi-mCherry more strongly than do Abi-mCherry and E-cadherin. One-way ANOVA (F(3,49)=104.0, p<0.0001 with post-hoc Tukey’s test; ****p<0.0001. **G**, Images showing the location of puncta of Fat2-3xGFP, Lar-3xGFP, or Sema5c-3xGFP along the length of individual phalloidin-stained filopodia. Abi-mCherry marks filopodia tips. **H**, Plots showing distributions of proteins in (G) along filopodia lengths. Fat2-3xGFP, Lar-3xGFP, Sema5c-3xGFP are all enriched with Abi-mCherry at filopodia tips. **I**, Diagram showing the organization of a leadingtrailing interface. Fat2, Lar, and Sema5c molecules reside together in puncta that span leading and trailing edges, with Fat2 enriched at the trailing edge and Lar and Sema5c at the leading edge.

Two biochemical signaling modules operate at leading-trailing interfaces, where they coordinate the migratory behaviors of neighboring follicle cells. The first module is composed of the atypical cadherin Fat2 and the receptor tyrosine phosphatase Leukocyte-antigen-related-like (Lar). Fat2 is enriched along the trailing edge of each cell, where it acts in trans to concentrate Lar and the WAVE complex across the cell-cell interface, at the leading edge of the cell behind^18–20^ (Fig. 1C,D). Lar also contributes to WAVE complex localization, but not as strongly as Fat2, implying the existence of additional unidentified Fat2 effectors^19,21^. Together, these proteins restrict cell protrusive activity to a single leading edge domain and orient all the cells’ protrusions in a uniform direction across the tissue^20^. The second module is composed of a transmembrane semaphorin (ligand) and plexin (receptor) pair, Semaphorin 5c (Sema5c) and Plexin A (PlexA), which are enriched at leading and trailing edges respectively^22^ (Fig. 1C,D). In other contexts, semaphorin-plexin signaling can lower integrin-based adhesion and/or inhibit protrusivity on the plexin-containing cell side^23–25^. Similarly, overexpression of Sema5c in one follicle cell reduces the protrusivity of its neighbors in a PlexA-dependent manner^22^. This led to the model that Sema5c signals through PlexA to maintain a non-protrusive state at cells’ trailing edges.

Despite their distinct depletion and overexpression phenotypes, several lines of evidence suggest that the Fat2-Lar and Sema5c-PlexA modules function within one interface-polarizing signaling system. A series of pairwise comparisons show that Fat2, Lar, and Sema5c all colocalize with the WAVE complex in interface-spanning puncta that sit at the tips of filopodia within a broader lamellipodium^19–22^ (Fig. 1E-I). For reasons that are not yet clear, PlexA only rarely colocalizes with the other proteins^22^. Loss of Lar also reduces Sema5c’s enrichment at leading edges, implying functional interaction between the two modules in addition to their shared spatial organization^22^. In this study, we investigated the hierarchy of interactions between Fat2, Lar, and Sema5c by which they form interface-spanning puncta, and asked how the three proteins work together to promote collective migration.

We found that Fat2 forms the core of both the Lar- and Sema5c-containing signaling modules. Trailing edge-enriched Fat2 concentrates both Lar and Sema5c at leading edges in trans and is continuously required for them to maintain this localization. Fat2 performs this function, at least in part, by slowing Lar’s and likely Sema5c’s turnover from the leading edge. We further found that Lar and Sema5c act in parallel to promote collective migration. From these data, we propose that Fat2 acts as a central organizer of follicle cells’ interface-polarizing signaling system, serving to couple and polarize the distributions of multiple effectors that together align the motility machinery of neighboring cells.

## Results

### Fat2 maintains both Lar and Sema5c’s enrichment at leading edges

Fat2, Lar, and Sema5c all colocalize in interface-spanning puncta, with Fat2 at trailing edges and Lar and Sema5c at leading edges^18,19,22^. Fat2 acts in trans to concentrate Lar at leading edges^19^, a result we confirmed in this study using a new endogenous 3xGFP tag on Lar (Lar-3xGFP, Figs. 2A,B; S1). To ask if Fat2 has the same effect on Sema5c, we generated epithelia with mosaic expression of *fat2*-RNAi and measured Sema5c-3xGFP levels along leading-trailing interfaces. Sema5c levels were reduced wherever a *fat2*-RNAi cell was present ahead of the interface, regardless of the genotype of the cell behind the interface (Fig. 2C,D), again indicating a local trans interaction. Since Lar also contributes to Sema5c’s enrichment at leading edges^22^, we asked if Fat2 localizes Sema5c indirectly through Lar. We confirmed that Sema5c levels were indeed reduced at leading-trailing interfaces between control cells and cells with a null *lar* allele (*lar*^*13*.*2*^, Fig. 2E,F). However, this reduction was not as great as at interfaces between *fat2*-RNAi cells (Fig. 2G). We conclude that Fat2 acts in trans to concentrate Sema5c at leading edges at least partly independently of Lar (Fig. 2H).

**Figure 2:**
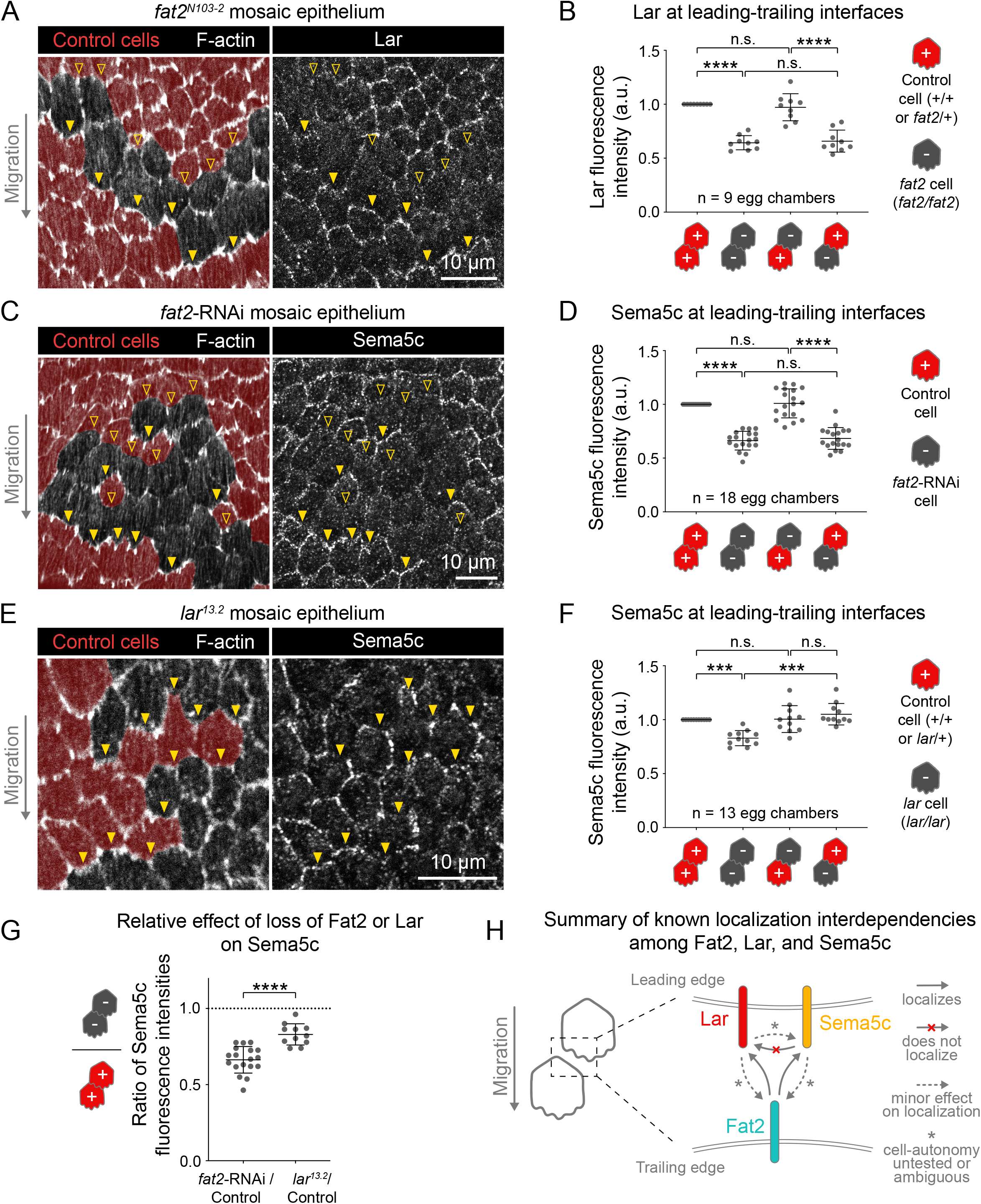
Fat2 concentrates Lar and Sema5c in trans at the leading edge. **A,C,E**, Images of genetically-mosaic epithelia expressing Lar-3xGFP or Sema5c-3xGFP. Control cells are pseudo-colored red based on genetically-encoded markers; homozygous mutant or RNAi-expressing cells are unlabeled. Phalloidin staining of actin protrusions indicates the migration direction. Arrows indicate leading-trailing interfaces at genotype boundaries, where the cell-autonomy localizing interactions can be assessed. Arrows are filled where Lar-3xGFP or Sema5c-3xGFP enrichment appears normal and hollow where it is reduced. **B,D,F**, Plots of fluorescence intensity at leading-trailing interfaces between pairs of cells of different genotype combinations. Bars indicate mean ± SD. **A**, Images of a *fat2*^*N103-2*^ mosaic epithelium expressing Lar-3xGFP. **B**, Plot of Lar-3xGFP in *fat2*^*N103-2*^ mosaic epithelia. Lar-3xGFP is reduced at the leading edge of cells of any genotype behind *fat2*^*N103-2*^ cells. Repeated measures one-way ANOVA (F(3, 24)=60.38, p<0.0001 with post-hoc Tukey’s test; n.s. (left to right) p=0.66, 0.96, ****p<0.0001. **C**, Images of a *fat2*-RNAi mosaic epithelium expressing Sema5c-3xGFP. **D**, Plot of Sema5c-3xGFP in *fat2*-RNAi mosaic epithelia. Sema5c-3xGFP enrichment is reduced at the leading edge of cells of any genotype behind *fat2*-RNAi cells. Repeated measures one-way ANOVA (F(2.42, 41.07)=96.23, p<0.0001 with post-hoc Tukey’s test; n.s. (left to right) p=0.99, 0.79, ****p<0.0001. **E**, Images of a *lar*^*13*.*2*^ mosaic epithelium expressing Sema5c-3xGFP. **F**, Plot of Sema5c-3xGFP in *lar*^*13*.*2*^ mosaic epithelia. Sema5c-3xGFP enrichment is reduced at leading-trailing interfaces between two *lar*^*13*.*2*^ cells, but not at interfaces with a control cell either ahead or behind, which is inconsistent with Lar simply recruiting Sema5c locally to their shared leading edge. Repeated measures one-way ANOVA (F(2.24, 22.41)=19.93, p<0.0001 with post-hoc Tukey’s test; n.s. (left to right) p>0.99, 0.77, ***p=0.003, ****p<0.0001. **G**, Plot comparing the effects of Fat2 loss and Lar loss on Sema5c-3xGFP enrichment at leading-trailing interfaces. Data replotted from (D) and (F). Both *fat2*-RNAi and *lar*^*13*.*2*^ reduce Sema5c-3xGFP enrichment (fluorescence intensity ratios below dotted line at *y*=1), but *fat2*-RNAi causes a greater reduction. Unpaired t-test; ****p<0.0001. **H**, Diagram showing dependencies among Fat2, Lar, and Sema5c for localization to the leading-trailing interface, summarizing findings from this and previous studies^19,22^ (see also Fig. S3). Fat2 concentrates Lar and Sema5c at the leading edge in trans, and Lar and Sema5c play minor, perhaps non-interface-local roles in the distribution of the three proteins.

We next asked whether Fat2 is continuously required for Lar and Sema5c’s enrichment at leading edges, or whether it becomes dispensable after interface polarity is established. Because calcium ions are integral to the structure of cadherin extracellular domains^26–28^, calcium removal should disrupt any Fat2 function that depends directly or indirectly on its extracellular domain. Treating egg chambers with the cell-impermeable calcium chelator ethylene glycol-O,O′-bis(2-aminoethyl)-N,N,N′,N′-tetraacetic acid (EGTA, 20 mM) for five minutes reduced the levels of protrusion-associated F-actin (Fig. 3A), consistent with loss of Fat2 function^19,21^ (compare to Fig. 2A,C). It did not, however, cause other obvious changes to cell morphology, and protrusions returned within one hour after EGTA washout (Fig. 3A), suggesting that this method rapidly and reversibly inhibits Fat2 without generally disrupting tissue architecture. Notably, the five minute EGTA treatment substantially reduced Lar and Sema5c levels along leading-trailing interfaces (Lar fluorescence intensity by 50%, Sema5c by 25%, Fig. 3B,C). Fat2 levels were nearly unchanged after five minutes, but its distribution became even more punctate, and by 30 minutes of EGTA treatment its levels had substantially decreased (Fig. 3B-D). Thus, Fat2 is continuously required to maintain the enrichment of Lar and Sema5c at leading edges.

**Figure 3:**
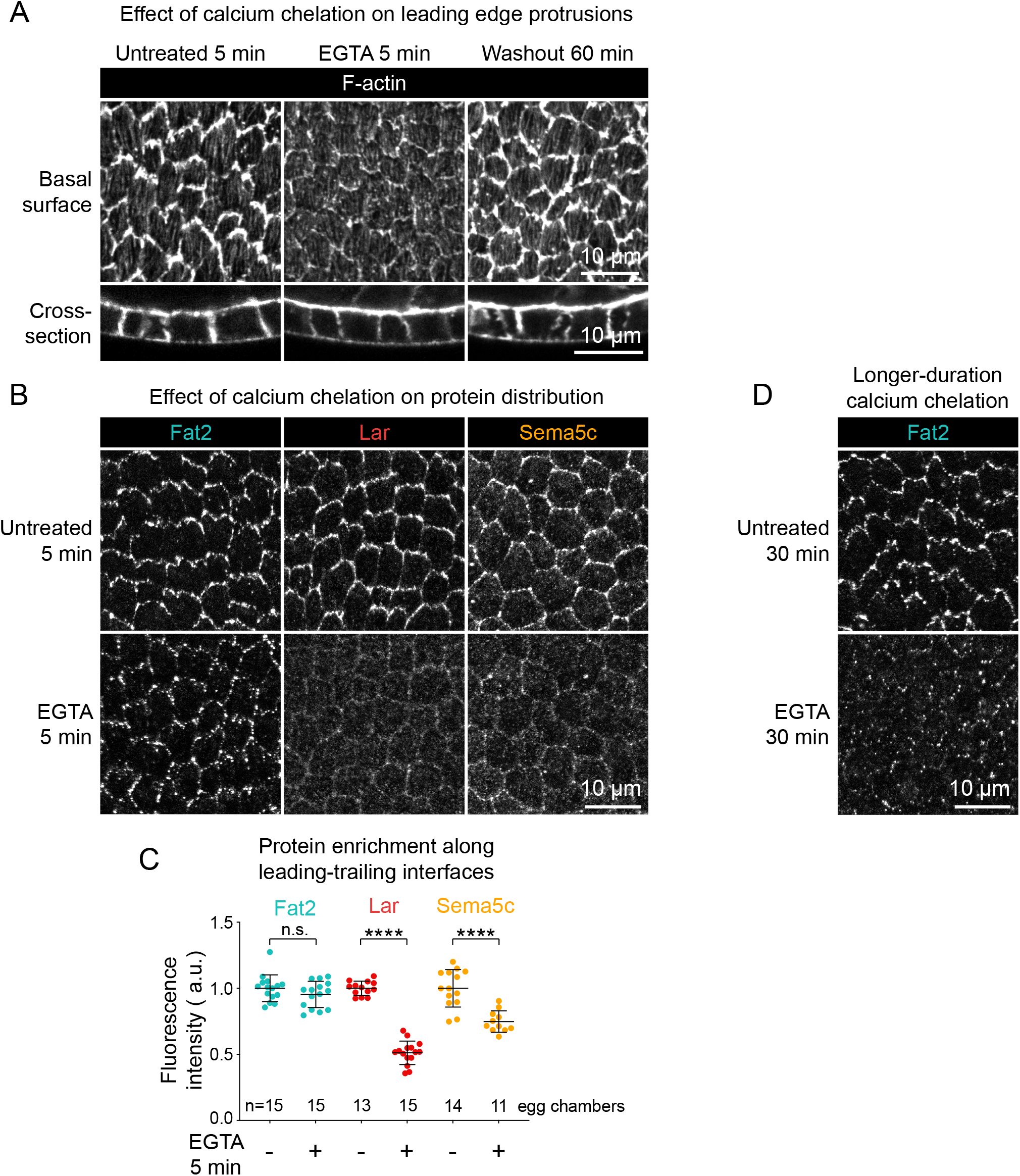
Calcium chelation causes rapid loss of Lar and Sema5c from the leading edge. **A**, Images of the basal surfaces of the follicle cells with F-actin stained by phalloidin after being left untreated for 5 minutes, treated with 20 mM EGTA (an extracellular calcium chelator) for 5 minutes, or treated with EGTA for 5 minutes followed by washout and 60 minutes of recovery. F-actin-rich protrusions are reduced by EGTA treatment, but return following washout. Cross-section images show no apparent changes to follicle cell shape or epithelial integrity caused by EGTA treatment or treatment and washout. **B**, Images of Fat2-, Lar-, or Sema5c-3xGFP at the basal surfaces of epithelia left untreated or treated with 20 mM EGTA for 5 minutes, representative of data quantified in (C). **C**, Plot of the mean fluorescence intensity of Fat2-3xGFP, Lar-3xGFP, and Sema5c-3xGFP along leading-trailing interfaces at the basal surface, with or without 5 minutes of EGTA treatment. EGTA treatment causes a reduction in Lar- and Sema5c-3xGFP levels, but no significant change in Fat2-3xGFP levels. Bars indicate mean ± SD. Unpaired t-tests; n.s. p=0.21, ****p<0.0001. **D**, Images of Fat2-3xGFP at the basal surface of epithelia left untreated or treated with 20 mM EGTA for 30 minutes. Unlike in (B) and (C), this longer treatment results in nearly-complete clearance of Fat2-3xGFP from the basal surface.

To probe the limits of Fat2’s capacity to position the leading-edge proteins, we used GrabFP-A^Int^ to create an ectopic Fat2 population^29^ (Fig. S2A), and asked if this could relocalize Lar. At present, we lack a functional Sema5c antibody, and so could not check for its relocalization. Fat2 and Lar normally colocalize in puncta at the basal surface and along tricellular junctions that span the apical-basal axis^30,31^. Expression of GrabFP-A^Int^ caused Fat2-3xGFP to accumulate all around adherens junctions as well, and increased Fat2-3xGFP levels overall (Fig. S2B-F). The ectopic Fat2 had no effect on the distribution of Lar, which remained restricted to tricellular junctions in the adherens junction plane despite Fat2’s presence all around cell perimeters (Fig. S2D-F). These data show that Fat2 is limited in its ability to recruit Lar, and that there must be some other feature(s) of basal cell-cell interfaces and tricellular junctions that are required for Lar’s enrichment and perhaps for puncta assembly in general.

### Lar and Sema5c play only a minor role in the localization of Fat2

Given Fat2’s importance for localizing Lar and Sema5c, we asked if they play a reciprocal role in localizing Fat2. To this end, we measured the average level of Fat2-3xGFP along all cell-cell interfaces at the basal surface of epithelia lacking Lar (*lar*-RNAi), Sema5c (*Sema5c*^*K175*^), or both proteins together (*lar*-RNAi, *Sema5c*^*K175*^) and compared these values to control epithelia. Full basal cell perimeters were used instead of leading-trailing interfaces because Fat2 loses its planar polarization when migration is disrupted^19^, as occurs in some of these backgrounds (see Fig. 5). Fat2 levels were normal in epithelia lacking Lar and mildly reduced in epithelia lacking Sema5c (Fig. S3A-C). Importantly, Fat2 levels were not further reduced in epithelia lacking both proteins compared to those lacking Sema5c alone (Fig. S3A,B). So, whereas Fat2 plays a major role in localizing Lar and Sema5c, Lar and Sema5c play at most a minor role in localizing Fat2 (Fig. 2H).

### Slowly-exchanging Fat2 molecules decrease the turnover of the leading edge proteins

We hypothesized that Fat2’s ability to concentrate Lar and Sema5c across their shared interface might be due to Fat2 slowing their turnover from cells’ leading edges. To test this idea, we first used fluorescence recovery after photobleaching (FRAP) to measure the turnover rates of all three proteins along leading-trailing interfaces at the basal surface of control epithelia. Within 6 minutes after photobleaching, Fat2-3xGFP fluorescence had only recovered to 29% ± 10% of its initial levels, whereas Lar-3xGFP and Sema5c-3xGFP fluorescence had recovered to 70% ± 21% and 90% ± 26%, respectively (Fig. 4A-E; Movie 1). During a longer recovery period of 28 minutes, Fat2-3xGFP fluorescence increased to 59% ± 25% of initial levels, still lower than the maximum recovery seen for Lar and Sema5c (Fig. 4F,G; Movie 2). These data show that Fat2 molecules reside at leading-trailing interfaces for tens of minutes, and Lar and Sema5c only minutes, under normal conditions.

**Figure 4:**
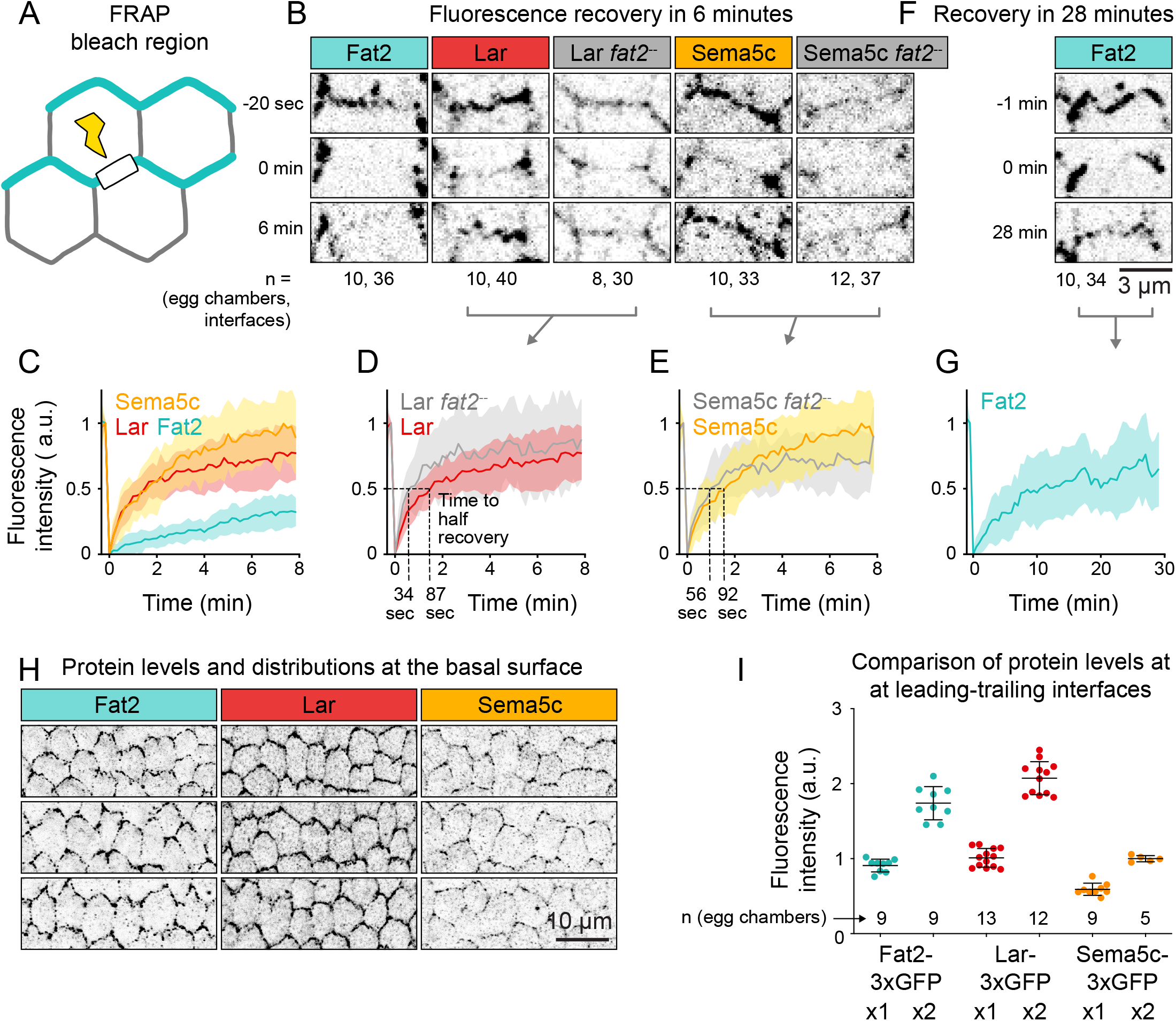
Fat2 is relatively stable at the trailing edge, and slows the departure of Lar and Sema5c from the leading edge. **A**, Diagram of the photobleached regions: leading-trailing interfaces at the basal surface, excluding the two bounding tricellular junctions. **B**, Example images of Fat2-3xGFP, Lar-3xGFP, and Sema5c-3xGFP at individual leading-trailing interfaces shortly before, immediately after, and 6 minutes after photobleaching. The *fat2*^*N103-2*^ were non-migratory and lacked planar polarity so any interfaces were used. The displayed regions follow interface movement. **C**, Plot comparing recovery of Fat2-3xGFP, Lar-3xGFP, and Sema5c-3xGFP fluorescence intensity to leading-trailing interfaces over 8 minutes following photobleaching. Solid lines and filled regions show mean ± SD. Fat2 recovers more slowly than Lar or Sema5c. **D**, Plot comparing recovery of Lar-3xGFP in control epithelia (reproduced from (C)) and *fat2*^*N103-2*^ epithelia. The times at which half the fluorescence had recovered are indicated for each. The speed at which Lar fluorescence recovers to half its initial level was more than doubled in the absence of Fat2. **E**, Plot comparing recovery of Sema5c-3xGFP in control epithelia (reproduced from (C)) and *fat2*^*N103-2*^ epithelia. The times at which half the fluorescence had recovered are indicated for each. The speed of Sema5c turnover may be slightly higher in the absence of Fat2. **F**, Images of Fat2-3xGFP at a leading-trailing interface shortly before, immediately after, and 28 minutes after photobleaching. The displayed regions follow interface movement. **G**, Plot of Fat2-3xGFP fluorescence recovery over 28 minutes. In that time, its fluorescence had still only recovered to 65% of its initial level, less than Lar-3xGFP or Sema5c-3xGFP did within 8 minutes. **H**, Images of Fat2-3xGFP, Lar-3xGFP, and Sema-5c-3xGFP at the basal surfaces of rows of cells from multiple epithelia, acquired and displayed using the same settings. **I**, Plot comparing the levels of Fat2-3xGFP, Lar-3xGFP, and Sema-5c-3xGFP at leading-trailing interfaces, expressed in two copies or with a wild-type allele. No correction for background fluorescence was performed. Interface enrichment differed across the three proteins, and Sema-5c-3xGFP levels were approximately half those of Fat2-3xGFP and Lar-3xGFP.

We then asked if Fat2 regulates the turnover of Lar and Sema5c at interfaces by performing the same FRAP experiment in epithelia lacking Fat2. Indeed, recovery of Lar-3xGFP fluorescence was faster in *fat2*^*N103-2*^ epithelia than in control epithelia (time to 1/2 recovery: 34 sec in *fat2*^*N103-2*^, 87 sec in control; Fig. 4B,D; Movie 1). The initial recovery of Sema5c-3xGFP was also slightly faster on average in the absence of Fat2 (time to 1/2 recovery: 56 sec in *fat2*^*N103-2*^, 92 sec in control; Fig. 4B,E; Movie 1), but the trend differed during later recovery, complicating interpretation of this data. We note that Sema5c-3xGFP levels at interfaces were lower than those of Lar-3xGFP normally (Fig. 4H,I), and were even lower in *fat2*^*N103-2*^ epithelia, which may have impaired our ability to resolve changes in recovery speed. Altogether, these data show that Fat2 has a stabilizing effect on Lar and perhaps on Sema5c, and suggest that Fat2 concentrates them at leading edges by slowing their turnover rather than by speeding their arrival.

### Lar and Sema5c promote collective migration in parallel downstream of Fat2

Fat2, Lar, and Sema5c are all required for normal collective migration, but whereas loss of Fat2 prevents migration entirely, loss of Lar only slows migration, and loss of Sema5c both slows it and delays its onset^18,19,22^ (Fig. 5A,B). These milder migration phenotypes suggest that Lar and Sema5c may act in parallel downstream of Fat2 to regulate cell motility. If true, analysis of the cell-scale phenotypes caused by loss of Lar or Sema5c could help us understand the two “arms” of the interface signaling system that Fat2 assembles. Fat2 promotes epithelial migration both by increasing cell protrusivity and by polarizing cells’ protrusions in the direction of tissue movement^19–21^, so we asked if Lar or Sema5c play similar roles in protrusion formation and polarity. These roles have previously been studied using F-actin labeling^19,21,22^, but we recently found that this approach misses much of the protrusive activity in *fat2*^*N103-2*^ epithelia^20^. We therefore used live imaging and membrane labeling to re-examine the protrusion phenotypes caused by loss of Lar or Sema5c alone, and to build on this by determining the effect of losing both proteins together.

First, we examined Lar’s role in protrusion formation and polarity. F-actin staining had suggested that cells lacking Lar are less protrusive than normal^19,21^. However, using membrane labeling, we detected no reduction in protrusivity in *lar*^*13*.*2/bola1*^ epithelia (Fig. 5C,D; Movie 3). This suggests that Lar increases F-actin enrichment within protrusions, but is not required for protrusion formation. Loss of Lar did, however, disrupt protrusion polarity. Whereas control epithelia had most protrusions pointed in the direction of migration, epithelia lacking Lar had a sizeable minority pointed rearwards, and their protrusions were less aligned with migration overall (Figs. 5C,E; S4; Movie 3). These data show that Lar helps polarize protrusions across the tissue, and suggest that Fat2’s role in this process is due in part to its ability to localize Lar.

We then performed the same analysis for Sema5c. *Sema5c*^*K175*^ epithelia initiate migration later in development than normal, and their migration is slow and difficult to preserve ex vivo^22^. Consistently, about 1/2 of the *Sema5c*^*K175*^ epithelia were migrating detectably, albeit slowly, and several others had a degree of protrusion polarity that likely indicated they had been migrating prior to dissection Figs. 5A,B; S4B). In agreement with previous findings based on F-actin staining^22^, we did not observe a difference in the membrane protrusivity of *Sema5c*^*K175*^ epithelia. In the detectably-migrating subset of epithelia, protrusion polarity also appeared normal (Figs. 5C-E; S4; Movie 3). The slow migration speeds of these epithelia therefore do not seem to be caused by protrusion defects, although it is possible that Sema5c regulates protrusions in ways we did not detect. Alternatively, Sema5c may primarily regulate other aspects of cell motility such as contractility or adhesion.

Finally, we asked whether Lar and Sema5c work in parallel to promote collective migration by expressing *lar*-RNAi in follicle cells of *Sema5c*^*K175*^ egg chambers. These epithelia failed to migrate and had unpolarized protrusions in all cases, making them indistinguishable from *fat2*^*N103-2*^ epithelia (Figs. 5A-C,E; S4; Movie 3). We conclude that Lar and Sema5c act in parallel to regulate cell motility, likely through distinct effectors. These data further suggest that Fat2’s polarization of both Lar and Sema5c to leading edges is a major means by which it regulates the collective migration of follicle cells.

**Figure 5:**
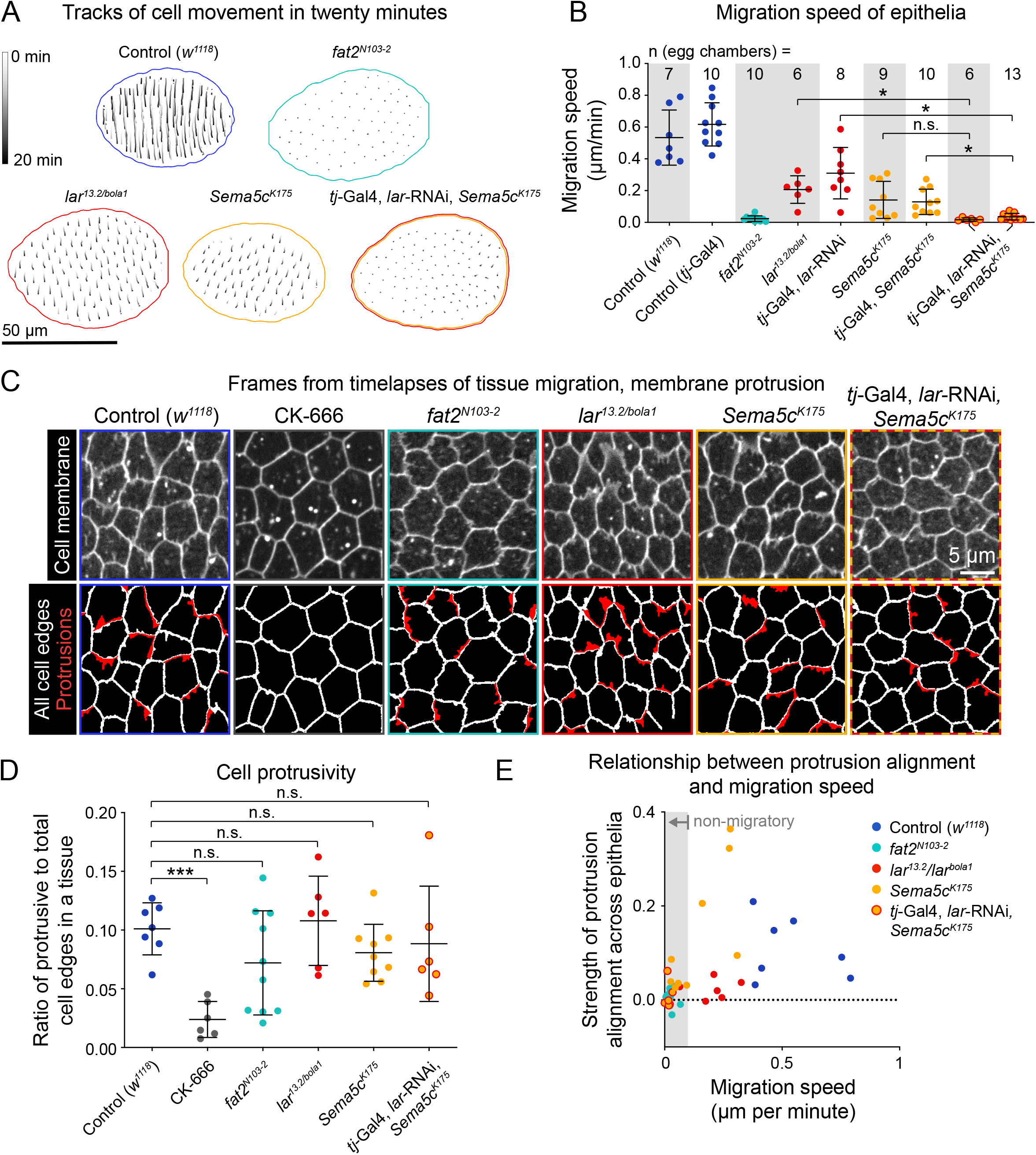
Lar and Sema5c act in parallel to polarize cell migratory machinery and promote collective migration. **A**, Tracks of cell movement over twenty minutes in individual epithelia. The in-focus tissue region is outlined. **B**, Plot of tissue migration speeds. Movies of conditions with white backgrounds on the graph were taken at medial planes through the follicle cells. Movies of conditions with gray backgrounds were taken at the basal surface and used for analysis of membrane protrusion traits in addition to migration speed. Whereas loss of Lar or Sema5c slows migration, simultaneous loss of both Lar and Sema5c stops it entirely. Gray backgrounds: Welch’s ANOVA (W(3.000,14.31)=64.11, p<0.0001) with post-hoc Dunnet’s T3 multiple comparisons test; *p=0.021, n.s. p=0.099. White backgrounds: Welch’s ANOVA (W(4.00,14.30)=21.64, p<0.0001) with post-hoc Dunnet’s T3 multiple comparisons test; * (left to right) p=0.0.011, n.s. p=0.025. **C**, Frames from timelapse movies used for measurement of migration speed and protrusion traits. Control epithelia treated with CK-666 are used as a non-protrusive control. The top row shows cell membranes labeled with CellMask. The bottom row shows the segmented cell edges and protrusions. Edges categorized as protrusive based on their average lengths of extension are shown in red. **D**, Plot of the level of cell protrusivity per epithelium, defined as the ratio of protrusive to total cell edges. Edges are categorized as protrusive if their average length is greater than that of 98% of edges in CK-666-treated epithelia. Unlike CK-666 treatment, loss of Fat2, Lar, or Sema5c causes only minor changes to the level of cell protrusivity, if any. Welch’s ANOVA (W(5.000,16.43)=12.83, p<0.0001) with post-hoc Dunnet’s T3 multiple comparisons test; ***p=0.0002, n.s. (left to right) p=0.69, >0.99, 0.70, >0.99. **E**, Plot showing the relationship between migration speed and the strength of protrusion alignment. An alignment of one indicates perfect alignment of all protrusions, and zero indicates either random protrusion orientations or symmetrical protrusion in two opposite directions. The gray region indicates non-migratory epithelia (speed < 0.1 um/min). Protrusions of slowly-migrating *Sema5c*^*K175*^ epithelia (but not *lar*^*13*.*2/bola1*^ epithelia) are poorly aligned with one another.

## Discussion

The follicle cells use biochemical signaling across their leading-trailing interfaces to polarize their migration machinery at interface, cell, and tissue scales. Here we have shown that Fat2 is a central organizer of this signaling system. Fat2 acts at the trailing edge of each cell to concentrate both Lar and Sema5c at the leading edge of the cell behind, and it appears to do so, at least in part, by slowing their turnover at this site. By contrast, Lar and Sema5c play at most minor roles in the localization of Fat2. In this way, Fat2 coordinates the activities of two effector proteins with distinct functions, allowing them to work synergistically to promote highly persistent collective migration.

One defining feature of this Fat2-based signaling system is that most of the component proteins colocalize in interface-spanning puncta. Cadherins often self-organize into clusters^32–35^, making it likely that this punctate organization stems from Fat2. However, whether Fat2 recruits Lar and Sema5c into the puncta through direct binding or through intermediary proteins is unknown. An ectopic Fat2 pool did not cause redistribution of Lar, suggesting that additional inputs are necessary for recruitment. We also do not know how Sema5c’s receptor, PlexA, fits into this model. Like Fat2, PlexA is enriched at cells’ trailing edges and helps to localize Sema5c, and yet antibody staining indicates that PlexA only partially colocalizes with the Fat2-based puncta. The biggest open question is how Fat2 becomes localized to the trailing edge, as this appears to be the key event that polarizes the entire signaling system. Mechanical feedback from collective migration itself is required for Fat2’s polarization to trailing edges^19^, but the nature of this feedback, and whether Fat2 has a trans binding partner that further stabilizes its localization, will be important areas for future investigation.

This work also sheds light on how Lar and Sema5c work together to promote collective migration. Loss of either protein alone impairs migration but does not stop it^19,22^. By contrast, removing both proteins together fully blocks migration in a way that is indistinguishable from loss of Fat2. These data suggest that once Fat2 concentrates Lar and Sema5c to cells’ leading edges, they then act in parallel to promote collective migration, likely by polarizing distinct aspects of the migration machinery.

What aspects of the migration machinery do Lar and Sema5c each control? In the case of Lar, it seems to be part of the bridge between Fat2 and WAVE complex-dependent protrusions—both Fat2 and Lar increase protrusive F-actin enrichment at leading edges (Fat2 in trans and Lar in cis), and both also help polarize protrusions in the direction of migration. In addition, Lar acts in trans to promote retraction of the trailing edge of the cell ahead^19^, but the mechanistic basis for this trans function, and the degree to which it is separable from Lar’s cis function, remain undetermined. In the case of Sema5c, cell-scale loss-of-function phenotypes have proven more elusive, but enrichment of PlexA at trailing edges and the ability of overexpressed Sema5c to suppress protrusion in trans both point to the trailing edge as the likely site of regulation^22^.

By positioning both Lar and Sema5c, Fat2 integrates two features of interface signaling systems known to operate in other collectively migrating cell types but not yet seen together. The first feature is the use of a trailing edge-associated mechanical cue to orient protrusions in the cell behind^36,37^. Mechanical localization of Fat2 to trailing edges polarizes protrusions in the following cell, in part by localizing Lar^19^. The second feature is the use of contact inhibition of locomotion^38–40^, which causes cells to polarize away from one another by suppressing protrusion and/or increasing contractility at the point of contact^41,42^. By localizing Lar and Sema5c to leading edges, Fat2 positions them to enforce trailing edge behavior at the contacting edge of the cell ahead. Therefore, Fat2 translates a mechanical cue into bi-directional signaling across leading-trailing interfaces to coordinate cell migratory behaviors for collective migration.

## Methods and Materials

### Materials, data, and code availability

Code used in data analysis, all numerical datasets, and plasmid maps are available at https://github.com/a9w/ epithelial_migration_signaling. Images and movies are available upon request, and will be added to a public repository once the manuscript is finalized.

### *Drosophila* sources, care, and genetics

The sources and references of all *Drosophila melanogaster* stocks used in this study are listed in Supp. Table 1. and the genotypes of the lines corresponding to each figure panel are listed in Supp. Table 2. *Drosophila* were raised on a diet of cornmeal molasses agar food at 25^°^C. Experimental females were collected 0-3 days post-eclosion and aged in the presence of males with a diet of the same cornmeal molasses agar supplemented with finely powdered active dry yeast before dissection. In most experiments, they were aged for two to three days at 25^°^C. Higher temperatures were used to increase the expression of Gal4-driven transgenes in some cases, and longer aging times (up to 5 days) were used to obtain more FRT recombination events in some mosaic epithelia. The temperatures and yeasting times used in each experiment are listed in Supp. Table 3. Epithelia mosaic for RNAi expression were generated using the Flp-out system with Gal4 expressed under control of a heat shock promoter. To heat shock, experimental females were aged while undergoing the following 12 hr temperature cycle: 1 hr 37^°^C, 1 hr 25^°^C, 1 hr 37^°^C, 9 hr 25^°^C. After 5 cycles (2.5 days), they were then transferred to 29^°^C and given fresh yeast for an additional 12 hrs prior to dissection to increase RNAi expression strength and uniformity across clones.

### Generation of the Lar-3xGFP line

Endogenous Lar was tagged C-terminally with three enhanced GFP proteins in tandem separated by short linker sequences (3xGFP) using CRISPR following the general approaches described in Gratz et al. (2013, 2014)^43,44^. The same 3xGFP sequence had been used to generate Fat2-3xGFP and Sema5c-3xGFP proteins in previous studies^19,22^. The guide RNA target sequence 5’-GCTTCTGCTATCGCGCTGCACTGG-3’ was chosen with fly-CRISPR Target Finder^44^. The underlined sequence was cloned into the pU6-BbsI-chiRNA plasmid, the upstream G added for efficient U6-driven expression, and the bold sequence is the adjacent PAM motif. The homologous recombination donor plasmid contained homology arms approximately 2 kb in length flanking the insertion target site, acquired by amplification of genomic DNA from the y1 Mnos-Cas9.PZH-2A w* (nanos-Cas9) background^45^. The 3xGFP sequence was amplified from the pDsRed-Fat2-3xGFP plasmid^19^. A linker with a sequence encoding the amino acids GSGGSTVPRARDPPVAT connects the Lar C-terminus with the N-terminus of 3xGFP. Homology arms, linker, and 3xGFP DNA fragments were inserted into donor plasmid pDsRed-attP, which contains 3xP3-DsRed flanked by loxP sites for insertion screening and subsequent removal^44^. The linker-3xGFP insertion was made immediately before the Lar stop codon. Guide and homologous recombination plasmids were injected by Genetivision in the nanos-Cas9 background, providing a germline Cas9 source. F1 males were screened for 3xP3-DsRed, which is expressed in the eye, and then 3xP3-DsRed was excised by crossing to Cre-expressing flies (MKRS hsFLP/TM6b Cre). Successful insertion was confirmed with genomic DNA sequencing of the lar C-terminus and 3xGFP insertion region following 3xP3-DsRed excision. This insertion strategy leaves a loxP “scar” downstream of the *lar* protein-coding region.

### Egg chamber dissection

Ovaries were dissected into 500 *µ*L of live imaging media (Schneider’s *Drosophila* medium with 15% fetal bovine serum and 200 *µ*g/mL insulin) in a spot plate using one set of Dumont #55 forceps in the experimenters’ dominant hand and 1 set of Dumont #5 forceps in the non-dominant hand. A more detailed description and images of this dissection protocol are available in Cetera et al. (2016)^46^. Ovarioles were removed from both the ovary and their individual muscle sheaths by pinching the germarium with forceps and pulling anteriorly. For fixed imaging, egg chambers older than stage 9 were removed prior to fixation. For live imaging, egg chambers older than the egg chamber to be imaged were removed from the ovariole strands by slicing through the adjoining stalk cells with a 27-gauge hypodermic needle. Removal of older egg chambers results in more compression of the younger egg chambers between the slide and the coverslip.

### Live imaging sample preparation

Immediately after dissection, ovarioles were transferred to a fresh well of live imaging media using a p10 pipettor. For samples with membrane staining, CellMask Orange (Thermo Fisher Scientific, Waltham, MA, 1:500) was added and samples incubated for 15 minutes, and then samples were washed in live imaging media to remove excess dye before mounting. For samples treated with CK-666, following plasma membrane staining, ovarioles were instead transferred to live imaging media with 750 μM CK-666 (Millipore Sigma, St. Louis, MO) and then mounted immediately after in the same CK-666-containing media. Samples were mounted along with 51 μm beads on standard glass slides, and covered with 22×22 mm #1.5 coverslips. The beads support the coverslip, controlling the amount of tissue compression. Some compression enables imaging of the basal surface of a field of cells in a single plane, but too much interferes with the migration process. Coverslip edges were sealed with petroleum jelly to prevent evaporation during imaging. Samples were checked for damage using the membrane stain or other available fluorescent markers as indicators, and excluded if damage was present. Slides were imaged for one hour or less.

### EGTA treatment

Egg chambers were dissected into live imaging media, and then half were transferred into spot plate wells with 250 *µ*L of live imaging media (the “control” treatment), and the other half into 250 *µ*L of live imaging media containing 20 mM EGTA (Thermo Fisher Scientific). They were incubated for 5 or 30 minutes, depending on the experiment, with wells covered by a slide to slow evaporation, and then fixed and stained with phalloidin as detailed below.

### Immunostaining and F-actin staining

After dissection, ovarioles were fixed at room temperature for 15 minutes in 4% EM-grade formaldehyde in PBT (phosphate-buffered saline with 0.1% Triton X-100 detergent), and then washed for 3×5 minutes in PBT. For antibody staining, egg chambers were incubated in primary antibody (mouse anti-Lar monoclonal, 1:200, or mouse anti-Dlg monoclonal, 1:20) overnight at 4^°^C, washed 3×5 minutes in PBT, and then incubated in secondary antibody (Donkey anti-mouse polyclonal, 1:200) for 2-3 hours at room temperature. For phalloidin staining, samples were incubated in TRITC phalloidin (Millipore Sigma, 1:250) for 15 minutes at room temperature, or with 647 AlexaFluor phalloidin (Thermo Fisher Scientific, 1:100) for 2-3 hours at room temperature. For a more detailed phalloidin staining protocol, see Anderson et al. (2023)^47^. Phalloidin and secondary antibody incubations were performed concurrently where applicable. Samples were then washed for 3×5 minutes in PBT and mounted on a slide in a drop (∼40 *µ*L) of SlowFade Diamond antifade (Invitrogen), covered in a 22×50 mm #1.5 coverslip sealed with nail polish, and stored at 4^°^C until imaged. This mounting strategy results in compression of Stage 6-7 egg chambers, allowing imaging of the basal surfaces of many cells in a single plane.

### Microscopy

All micrograph images collected are of egg chambers at either stage 6 or 7, as assessed by their size and by the size and shape of their oocyte in cross-section. Images were acquired using a Zeiss LSM800 upright laser scanning confocal microscope with either a 40x/1.3 NA EC 386 Plan-NEOFLUAR or 63x/1.4 NA Plan-APOCHROMAT oil immersion objective. The system was controlled with Zen 2.3 Blue acquisition software (Zeiss). Imaging was performed at room temperature. All images and measurements are of a single Z-slice of the follicle cells’ basal surfaces unless otherwise noted. Exceptions are Fig. S2B and the bottom rows of Figs. 3A and S1A, which show cross-sections through the apical-basal axis; Fig. S2D, which show a near-apical plane (the plane of the adherens junctions); and the timelapse movies used to measure the migration speeds plotted in Fig. 5B on a white background, which were taken at a medial plane through the cells with respect to their apical-basal axis. Unless noted, imaging settings were held constant for all images of the same fluorescent protein shown in the same image panel and any corresponding plot panels, but settings affecting image brightness were optimized separately for each fluorophore when multiple are present in the same panel, or for data shown in separate panels. The exceptions are in Fig. 2C,D and 2E,F, which were collected with the same imaging settings to enable the comparison in 2G, and Fig. 4H,I, in which Fat2-3xGFP, Lar-3xGFP, and Sema5c-3xGFP were imaged and displayed in the same manner to enable comparison of their levels.

### Cell tracking and migration speed measurement

Twenty minute-long timelapse movies were acquired of stage 6-7 egg chambers stained with CellMask membrane dye. These were taken at either the medial epithelial plane (used for migration speed measurement only, conditions with white background in Fig. 5B) or the basal plane (for migration speed and protrusion trait measurement, conditions with gray backgrounds). In either case, cells were segmented in each frame using the pretrained “cytoplasm” model in Cellpose^48^. Subsequent analysis steps were performed in Python using scikit-image and scipy libraries^49,50^. The gaps between segmented cells were closed with a watershed algorithm seeded by the initial segmented regions. Segmented cells were then tracked by linking regions of high overlap in consecutive frames. Errors in segmentation and tracking were corrected manually using napari^51^. Migration speed was measured from a 22 *µ*m-wide band of cells around the egg chamber’s “equator,” half-way between anterior and posterior poles, where linear cell migration speed is fastest in migratory epithelia. The displacement vector of the centroid of each cell present in consecutive pairs of frames was calculated, and these vectors averaged to obtain a vector whose length and direction were used to determine the tissue migration speed and direction.

### Measurement of membrane protrusivity and protrusion alignment with migration

Protrusions were identified from timelapse movies of epithelia stained with CellMask membrane dye, which yield rich information about protrusion characteristics including their sizes and distributions, and often allow detection of more protrusive activity than is detectable by F-actin labeling. Protrusions were segmented using a watershed-based approach, and their average lengths and orientations measured, as described in Williams et al. (2022)^20^ and briefly here. Protrusion segmentation and trait measurement were performed in Python using scikit-image and scipy libraries. Following cell segmentation, the region of high membrane fluorescence at the interface between each pair of neighboring cells was segmented. This region includes the cell-cell interface and any protrusions that extend across it from either of the neighboring cells. We then approximated the cell-cell interface position running through this region by drawing the shortest path through the region that connects its bounding vertices. This divides the region into two portions belonging to each of the neighboring cells, which we call “edges.”

To obtain a benchmark for the width of non-protrusive cell edges, we identified the average width (area divided by interface length) of edges from CK-666-treated epithelia, which are nearly non-protrusive. Edges from all conditions were considered protrusive if their width (corresponding with their average length of membrane extension from the interface) was greater than that of 98% of those of CK-666-treated epithelia. We then calculated the average cell protrusivity of each epithelium, defined as its ratio of protrusive to total edges.

To measure protrusion polarity, we took the protrusive edges (“protrusions”), identified their “tip” and “base” as in Williams et al. (2022)^20^, and then assigned their orientation as the direction of the vector from base to tip. As a metric for protrusion alignment across an epithelium, we found the dot product of each pair of protrusions present in the same frame, and then took the mean of all dot products. For each protrusion pair, the dot product will be 1 if they have the same orientation, 0 if they are orthogonal to one another, and –1 if they have opposite orientations. Averaging these for an epithelium would yield an alignment score of 1 if all protrusions were perfectly aligned (high vectorial polarity), but could have a score of zero either if protrusions were randomly oriented (no polarity), or if half pointed in each direction along one axis (high axial polarity, no vectorial polarity). We report protrusions’ alignment with one another, rather than with the direction of migration, because this is still meaningful in non-migratory epithelia (whether truly non-migratory or only non-migratory ex vivo). In migrating epithelia, protrusion-protrusion alignment and protrusion-migration alignment measurements were highly correlated.

### Quantification of Fat2-3xGFP fluorescence at cell-cell interfaces and the basal surface

Egg chambers expressing Fat2-3xGFP were stained with anti-Dlg to label cell edges, stage 6-7 egg chambers were imaged, and the cells were segmented using the pretrained “cytoplasm” model in Cellpose applied to the Dlg channel. Subsequent analysis steps were performed in Python using scikit-image and scipy libraries. Gaps between segmented cells were closed with a watershed algorithm seeded by the initial segmented regions, and segmentation errors were then manually corrected using napari. Mean fluorescence intensity was calculated at cell-cell interfaces (boundaries between segmented cells dilated by 10 px) and the entire in-focus basal surface.

### Measurement of fluorescence intensity at leading-trailing interfaces

Leading-trailing interface regions were annotated by hand based on phalloidin staining using Fiji (ImageJ)^52,53^: 10 px-wide segmented lines were drawn along leading-trailing interfaces and then the mean fluorescence intensity across those interfaces was calculated. In experiments with non-mosaic epithelia, at least 10 leading-trailing interfaces were measured. For more information about measurement in mosaic epithelia, see the next section. Note that no correction was performed for background fluorescence, so the proportional relationship between fluorescence intensity measurements should not be interpreted as corresponding directly to the proportional relationship between protein levels.

### Analysis of localizing interactions and their cell autonomy

Analysis of protein localization in genetically mosaic epithelia was used to disentangle cell- and interface-scale protein-localizing interactions from the tissue-wide effects of genotype on planar polarity and migration. It was also used to determine the cell autonomy of localizing interactions. Interpretations of the cell autonomy of effects on Lar or Sema5c’s localization were made based on the much greater enrichment of Lar and Sema5c at leading edges than trailing edges in normal circumstances^19,22^ (see Fig. 1C). In epithelia mosaic for a loss-of-function mutation (generated using the Flp/FRT system), cells with the chromosome containing the wild-type allele were marked with a nucleus-enriched fluorophore, and “control” cells include both homozygous wild-type and heterozygous cells. In epithelia mosaic for RNAi expression (generated using the Flp-out system), RNAi-expressing cells were marked with a nucleus-enriched fluorophore. F-actin staining with AlexaFluor 647 phalloidin was used to visually assess whether epithelia were planar-polarized and migratory (based on stress fiber alignment across the epithelium), their migration direction (based on the orientation of leading edge protrusions), and to check for regions of tissue damage that could otherwise be misinterpreted as genetic clones. Egg chambers were included in subsequent analysis if they were within stage six to seven, had planar-polarized actin stress fibers, and had both control and mutant/RNAi-expressing cells in view with at least three leading-trailing interfaces between cells of each genotype combination. Mean fluorescence intensity at individual leading-trailing interfaces was then measured as described in the previous section, with all leading-trailing interfaces at genotype boundaries, and a similar number of clone-internal ones, included. Plots show intensities rescaled per sample such that the mean fluorescence intensity at leading-trailing interfaces between control cells is equal to 1, and were made using GraphPad Prism 9.

### Plotting of protein intensity along an “unrolled” cell perimeter

Plots of “unrolled” cell perimeters were used to display the distributions of Fat2, Lar, and GrabFP-AInt fluorescence around cell perimeters and the correspondence between those distributions. 10 px-wide segmented lines were drawn around the perimeter of cells in Fiji, and fluorescence intensities measured. Using Python, intensities were rescaled for ease of comparison and plotted. Specifically, Fat2-3xGFP and anti-Lar intensities were rescaled such that they had a range of 0 to 1 in the control dataset. GrabFP-AInt was rescaled proportionally to Fat2-3xGFP.

### Colocalization of proteins along the leading-trailing interface

Multi-channel confocal imaging was performed, and resulting images analyzed, to determine the degree of colocalization between pairs of fluorophore-tagged proteins (Abi-mCherry and Fat2-3xGFP, Lar-3xGFP, Sema5c-3xGFP, E-cadherin(Shg)-GFP) along leading trailing interfaces. Images of the basal plane of follicle cells were acquired with a 63x/1.4 NA Plan-APOCHROMAT oil immersion objective to minimize chromatic aberration. In Fiji, 10 px-wide segmented lines were drawn along rows of leading-trailing interfaces. At least 20 leading-trailing interfaces were included per egg chamber, and fluorescence intensities at each position along the lines measured. Spearman’s correlation coefficients were calculated for each egg chamber using the scipy statistics module in Python. As E-cadherin is fairly uniformly distributed along all cell-cell interfaces, correlation between E-cadherin-GFP and Abi-mCherry serves as a baseline for spurious correlation caused by the measurement region-drawing method and correlated effects of interface shape on the intensity distribution of both fluorophores. The plot was made using GraphPad Prism 9.

### Quantification of fluorescence distributions along filopodia

The fluorescence intensity distributions of proteins along the lengths of many filopodia were used to obtain representations of their typical distribution. Epithelia expressing Abi-mCherry and either Fat2-3xGFP, Lar-3xGFP, or Sema5c-3xGFP were stained with AlexaFluor 647 phalloidin. In Fiji, for one cell per egg chamber, 1 px-wide lines were drawn along the lengths of each filopodium identified phalloidin staining. Subsequent data processing and plotting was all performed in Python. Fluorescence intensity traces from along different filopodia were registered using the highest-intensity Abi-mCherry pixels, a proxy for the filopodia tips. Standard deviations were calculated for each fluorophore after rescaling traces so that their values each ranged from 0 to 1. For plotting, data were first averaged, then rescaled from 0 to 1.

### FRAP analysis of protein exchange at leading-trailing interfaces

Photobleaching was performed on samples mounted and imaged as described in previous sections using the timed bleaching tool in Zen Blue. Focusing at the basal surface plane, leading-trailing interface regions between two vertices, excluding those vertices, were selected for bleaching. Lar-3xGFP, *fat2*^*N103-2*^ and Sema5c-3xGFP, *fat2*^*N103-2*^ epithelia were non-migratory and lacked planar polarity, so any interfaces were used. Up to four interfaces were bleached per egg chamber, always from non-neighboring cells. Two frames were acquired before bleaching, bleaching performed once, and then timelapse acquisition continued for a total of either 8 minutes (at 10 sec/frame) or 30 minutes (at 30 sec/frame). Bleached interfaces were excluded from analysis if the majority of the interface length was not bleached, the bleached region encompassed a cell vertex, or the bleached interface was not in view throughout the timelapse. Fluorescence intensity measurements were made using Fiji. Rectangular regions were drawn around the bleached interfaces and the same number of non-bleached interfaces, the positions of these regions were translated in each time point as needed to follow interface movement, and the mean intensity of each was measured. Subsequent analysis and plotting was done in Python. Mean fluorescence per timepoint was calculated separately for bleached and unbleached regions, and the mean of the unbleached regions used to rescale the bleached regions to correct for the effects of progressive photobleaching and region selection inaccuracy. The bleach-corrected values were then rescaled so that the time point just before bleaching had a value of one, and the timepoint just after bleaching zero. Standard deviations were calculated for each timepoint from the mean intensities of individual bleached regions after they were rescaled in the same manner.

### Movie generation

For timelapse movies (but not analysis) of FRAP at cell-cell interfaces (Movies 1,2), bleach correction was performed on the full tissue timelapse using the “simple ratio” method in ImageJ prior to cropping and movie generation, and the cropped region was moved to follow the interface. For all movies, margins, scale bars, and text labels were first added to TIF image stacks using ImageJ or Adobe Illustrator 2021, and then the files exported from ImageJ as uncompressed AVI files. These were encoded as 1080 p30 MP4 files with H.264 (x264) video encoder using HandBrake 1.4.

### Reproducibility and statistical analysis

Two or more biological replicates were performed for each experiment, with each containing egg chambers pooled from multiple flies, and results confirmed to be qualitatively consistent between them. Visibly damaged egg chambers were excluded from all analyses. No randomization of treatment groups was performed, and experiments were not performed with blinding. Sample sizes were not predetermined using a statistical method. The number of egg chambers, interfaces, and/or filopodia (n) analyzed for each experiment can be found in the associated figure or figure legend. Statistical tests performed and their significance can also be found in figures or figure legends. An alpha of 0.05 was used to determine significance in all cases, but we have also attempted to plot data distributions in ways that allow the reader to weigh their similarities and differences for themselves. Statistical testing was all conducted using GraphPad Prism 9. Based on visual inspection, all datasets appeared normally distributed, and statistical tests assuming normalcy were used throughout. All t-tests were two-tailed. A one-way ANOVA was used when more than two conditions were compared. A repeated measures one-way ANOVA was used for analysis of genetic mosaic epithelia, in which multiple genetic conditions were present in the same tissue. Welch’s corrections were performed and Dunnet’s T3 multiple comparison tests used for datasets in which the variance did not appear consistent between conditions (indicated in figure legends). Otherwise, post-hoc Tukey’s multiple comparison tests were used. Initial and post-hoc multiple comparisons testing was conducted on all data present in the corresponding plots with two exceptions: in Fig. 3C, only data including the same fluorophore were compared, and in Fig. 5B, data from conditions with gray and white background shading were collected and analyzed separately.

## Author Contributions

A.M.W. and S.H.B. conceived of the study and contributed to experimental design. A.M.W. generated new reagents, performed experiments, analyzed data, and prepared figures. A.M.W. and S.H.B. wrote the manuscript.

## Acknowledgements

This study was supported by the National Institutes of Health (NIH) under grants R01 GM126047 and R35 GM148485 to S.H.B. and T32 HD055164 to A.M.W.. We thank members of the Horne-Badovinac and Munro Labs, Michael Glotzer, and Ellie Heckscher for discussion, and Seth Donoughe for image analysis advice.

## Supplemental Figures

**Figure S1:**
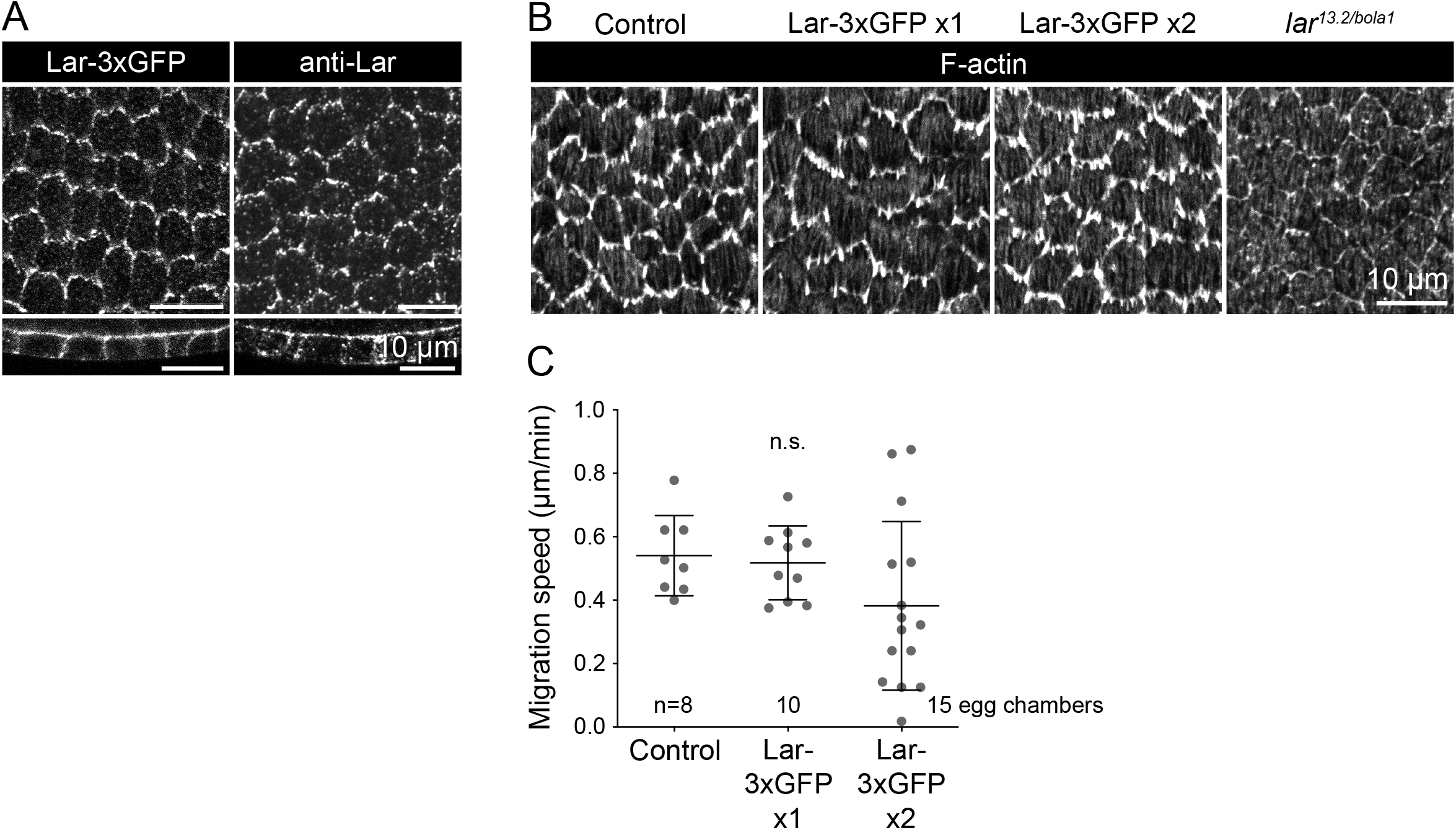
Lar remains largely functional with an endogenous 3xGFP tag. **A**, Images showing Lar distribution at the basal surface (top row) or in cross-section (bottom row) using either Lar-3xGFP or a Lar antibody. Lar localization patterns are similar between the two markers. **B**, Images showing F-actin (phalloidin) structures at the basal surface in epithelia with wild-type Lar, one or two Lar-3xGFP alleles, or with no Lar expression. F-actin protrusions appear normal in epithelia expressing Lar-3xGFP, in contrast to the reduced F-actin protrusions in epithelia without Lar. **C**, Plot of tissue migration speeds in control epithelia or epithelia with one or two Lar-3xGFP copies. Average migration speeds are similar to controls when one copy of Lar-3xGFP is present, but they are more variable and possibly slower in Lar-3xGFP homozygotes. Welch’s ANOVA (W(2.00,19.01)=1.92, p=0.17).

**Figure S2:**
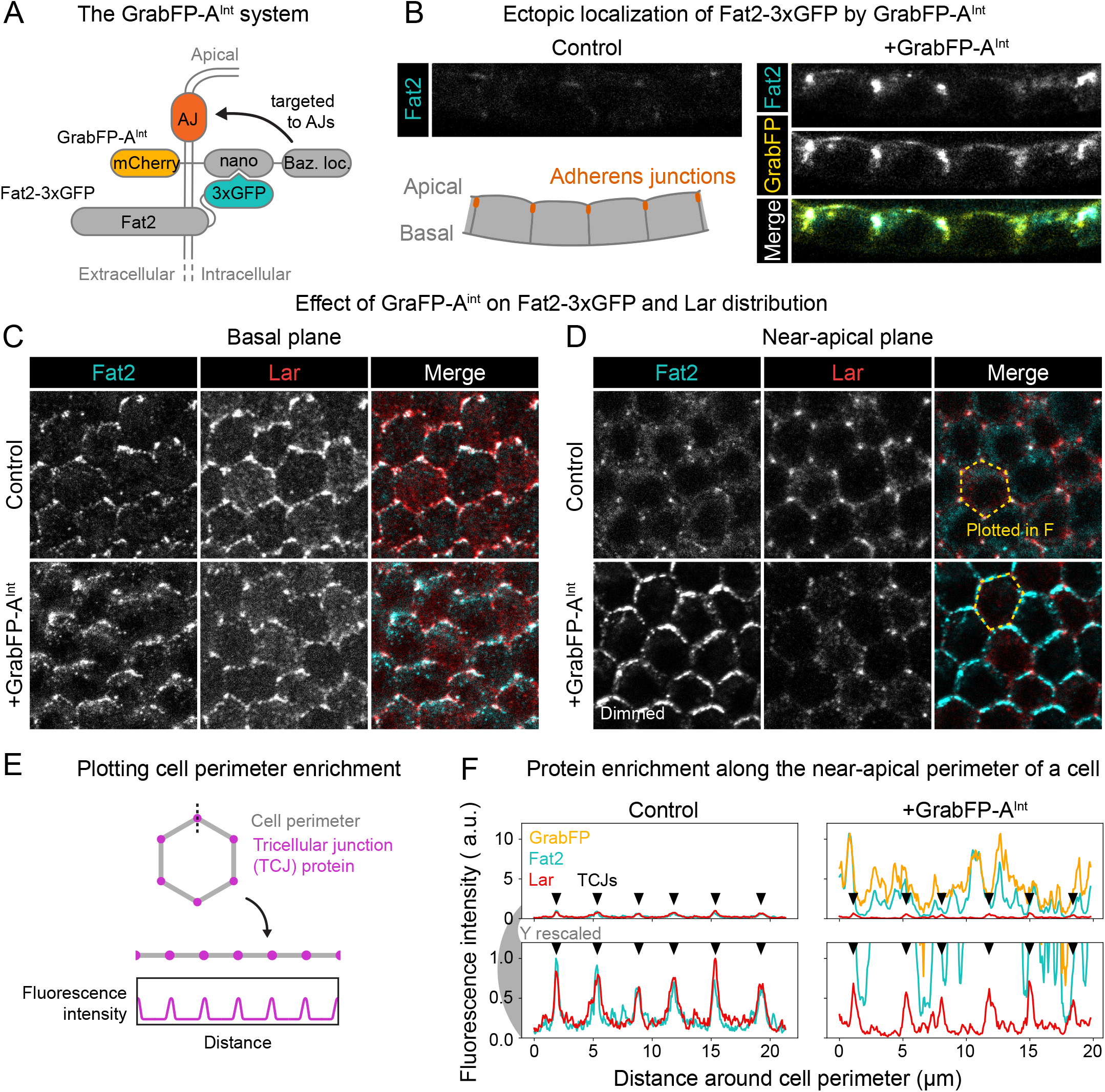
Fat2 is not sufficient to relocalize Lar away from the basal surface or tricellular junctions. **A**, Diagram showing how the GrabFP-A^Int^ protein is used to recruit Fat2-3xGFP to adherens junctions (AJs). GrabFP-A^Int^ binds intracellularly GFP-tagged proteins with a GFP nanobody (nano), and is targeted to AJs, along with any bound proteins, by a domain of Bazooka (Baz. loc.). **B**, Images of Fat2-3xGFP-expressing follicle cells in cross-section with or without expression of GrabFP-A^Int^. The diagram shows the imaging plane, with the adherens junctions to which GrabFP-A^Int^ is targeted indicated. Fat2-3xGFP is present in control cells, but its enrichment at adherens junctions and overall levels are strongly increased in the presence of GrabFP-A^Int^. **C**,**D**, Images of Fat2-3xGFP-expressing epithelia with or without co-expression of GrabFP-A^Int^, stained with a Lar antibody. Images in (C) show the basal planes of a group of cells; images in (D) show a near-apical plane through their adherens junctions. Fat2-3xGFP is displayed more dimly in the +GrabFP-A^Int^ image in (D) relative to other images to allow the Fat2-3xGFP distribution to be clear in each. **E**, Diagram illustrating the “unrolling’’ process by which the perimeters of cells such as those outlined in (D) are linearized for plotting in (F). **F**, Plots of the fluorescence intensity distribution of Fat2-3xGFP, anti-Lar, and GrabFP-A^Int^ (if present) along the cell perimeters outlined in (D). The upper row shows the full range of intensities, and the lower row shows an expanded view of the lower intensities. Triangles correspond to tricellular junctions (TCJs), to which Fat2 and Lar are normally largely restricted in this plane. GrabFP-A^Int^ expression causes a large increase in Fat2-3xGFP levels and their expansion around the entire cell perimeter, but Lar remains restricted to tricellular junctions.

**Figure S3:**
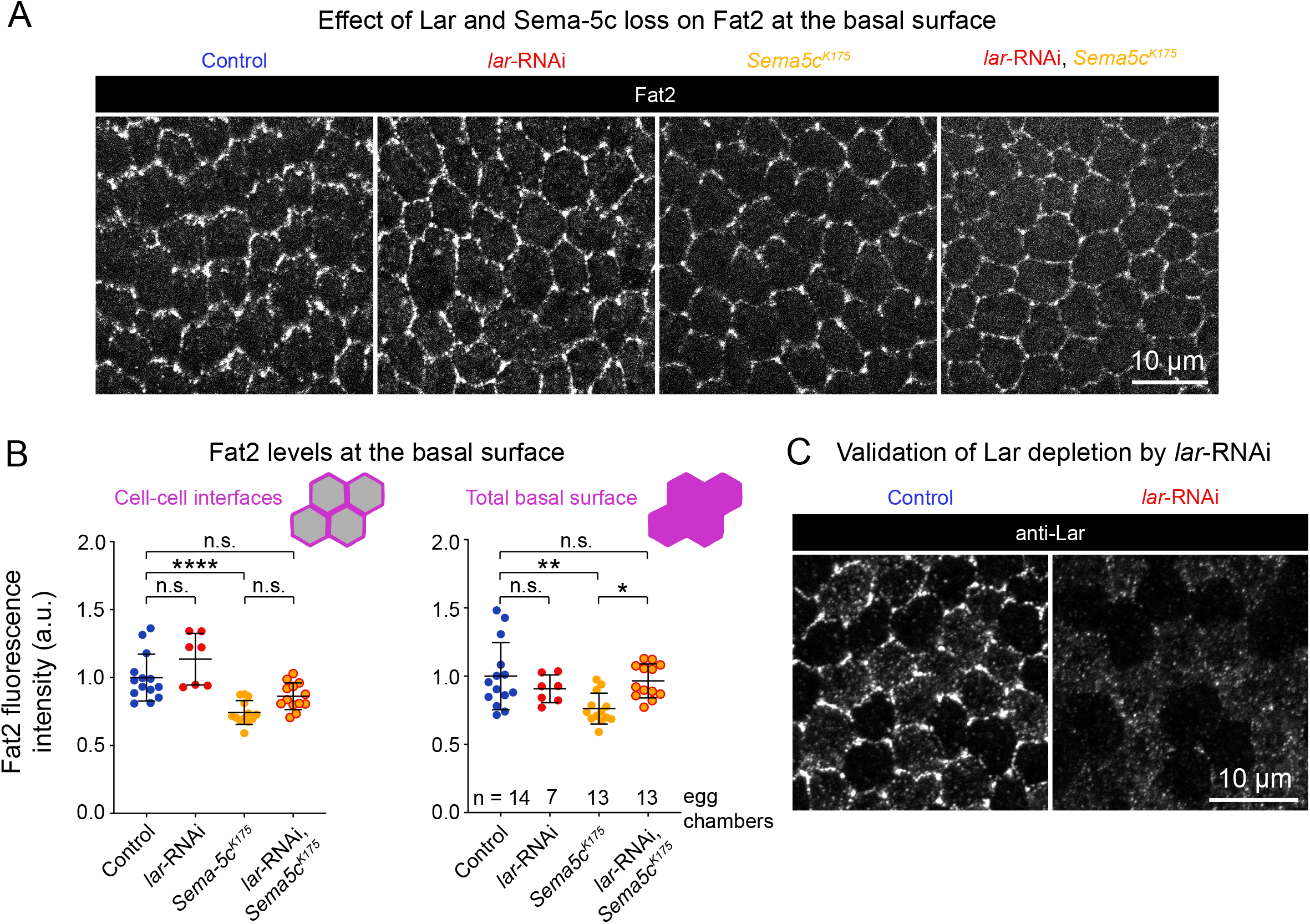
Fat2’s enrichment at interfaces shows only a mild dependence on Lar and Sema5c. **A**, Images of Fat2-3xGFP at the basal surface in control epithelia, *lar*-RNAi-expressing or *Sema5c*^*K175*^ epithelia, and in a combined *Sema5c*^*K175*^ *lar*-RNAi-expressing epithelium. **B**, Plots of Fat2-3xGFP levels at cell-cell interfaces (left) or the entire basal surface (right) of epithelia exemplified in (A). Fat2-3xGFP levels are slightly reduced at interfaces and across the basal surface in *Sema5c*^*K175*^ epithelia. Lar depletion causes no additional reduction in Fat2-3xGFP levels at these locations. Bars indicate mean±SD. Cell-cell interfaces: One-way ANOVA (F(3,43)=15.18, p<0.0001 with post-hoc Tukey’s test; ****p<0.0001, n.s. (left to right) p=0.16, 0.059, 0.14. Total basal surface: One-way ANOVA (F(3,43)=5.31, p=0.0033 with post-hoc Tukey’s test; **p=0.0030, *p=0.016, n.s. (left to right) p=0.63, 0.95. **C**, Images of Lar antibody staining at the basal surface, demonstrating the strength of Lar depletion by *lar*-RNAi.

**Figure S4:**
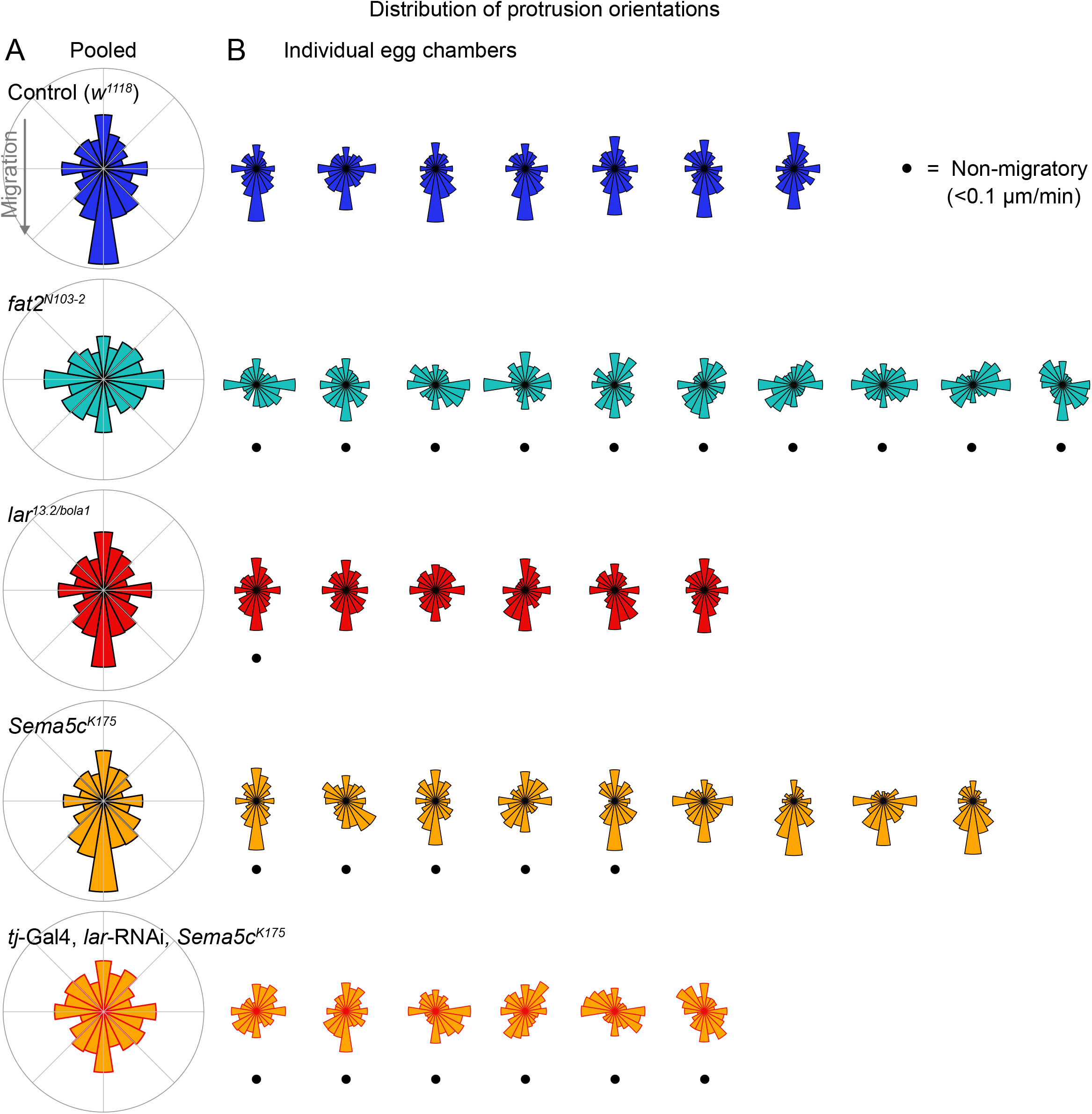
Loss of either Fat2, Lar, or Sema5c have different effects on the polarity of membrane protrusions. **A**,**B**, Polar histograms showing the distribution of protrusion orientations in epithelia pooled by genotype (A) or plotted individually (B). In (B), dots under histograms indicate that the epithelium was not migrating (speed <0.1 *µ*m/min). Anterior is left, posterior is right, and data has been flipped vertically as needed such that the migration direction (where applicable) is downwards. *lar*^*13*.*2/bola1*^ epithelia have excess rearwards protrusions, and removal of both Lar and Sema5c causes protrusions to be unpolarized, as is seen upon loss of Fat2.

## Movie captions

Movie 1: **FRAP of Fat2-3xGFP, Lar-3xGFP, and Sema5c-3xGFP at interfaces in control and *fat2***^***N103-2***^ **epithelia**. Movies begin 20 seconds before photobleaching and follow leading-trailing interfaces (control) or similarly-oriented interfaces (*fat2*^*N103-2*^) for the subsequent 8 minutes. Display brightness settings are adjusted separately for each fluorophore, but preserved between control and *fat2*^*N103-2*^ conditions. Associated with Fig. 4B-E.

Movie 2: **FRAP of Fat2-3xGFP at a leading-trailing interface over 30 minutes**. The timelapse movie follows fluorescence recovery at a leading-trailing interface beginning one minute before photobleaching. Associated with Fig. 4F,G.

Movie 3: **Effects of loss of Fat2, Lar, and Sema5c on migration and membrane protrusion dynamics**. Timelapse movies of the basal surfaces of epithelia labeled with CellMask membrane dye. Several examples from each condition are shown. Associated with Figs. 5C-E; S4.

**Supp.Table 1.**
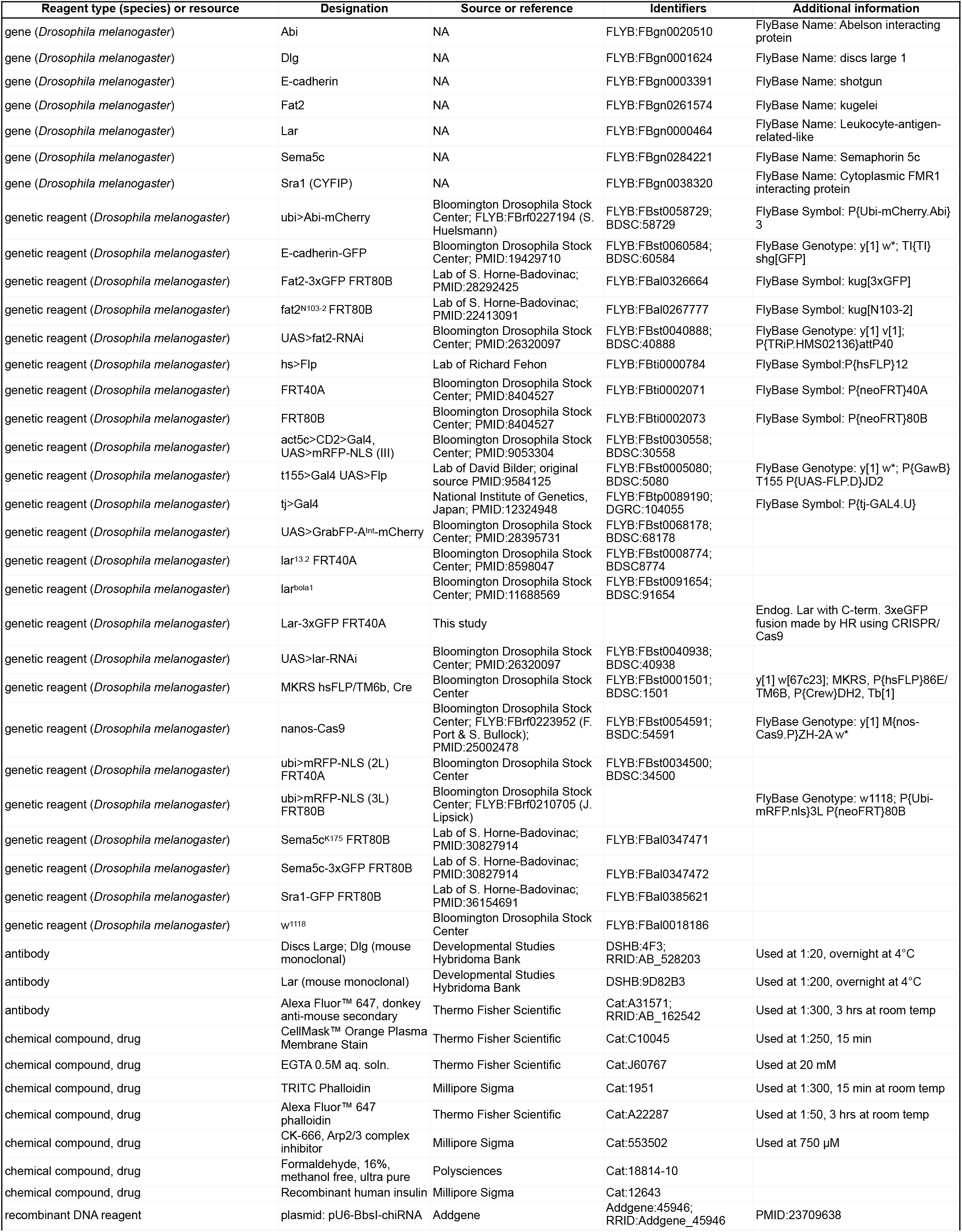
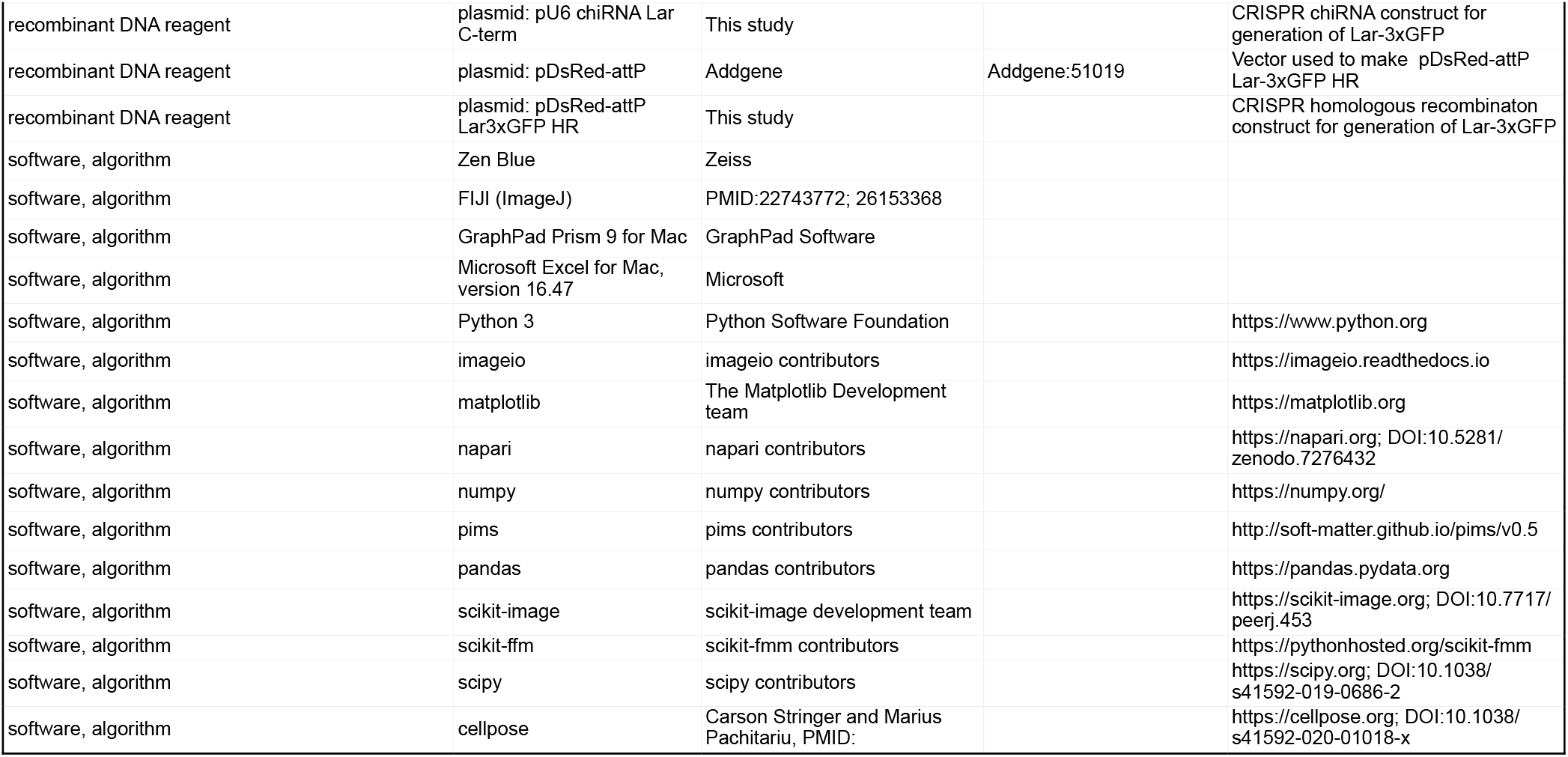
Key resources

**Supp.Table 2:**
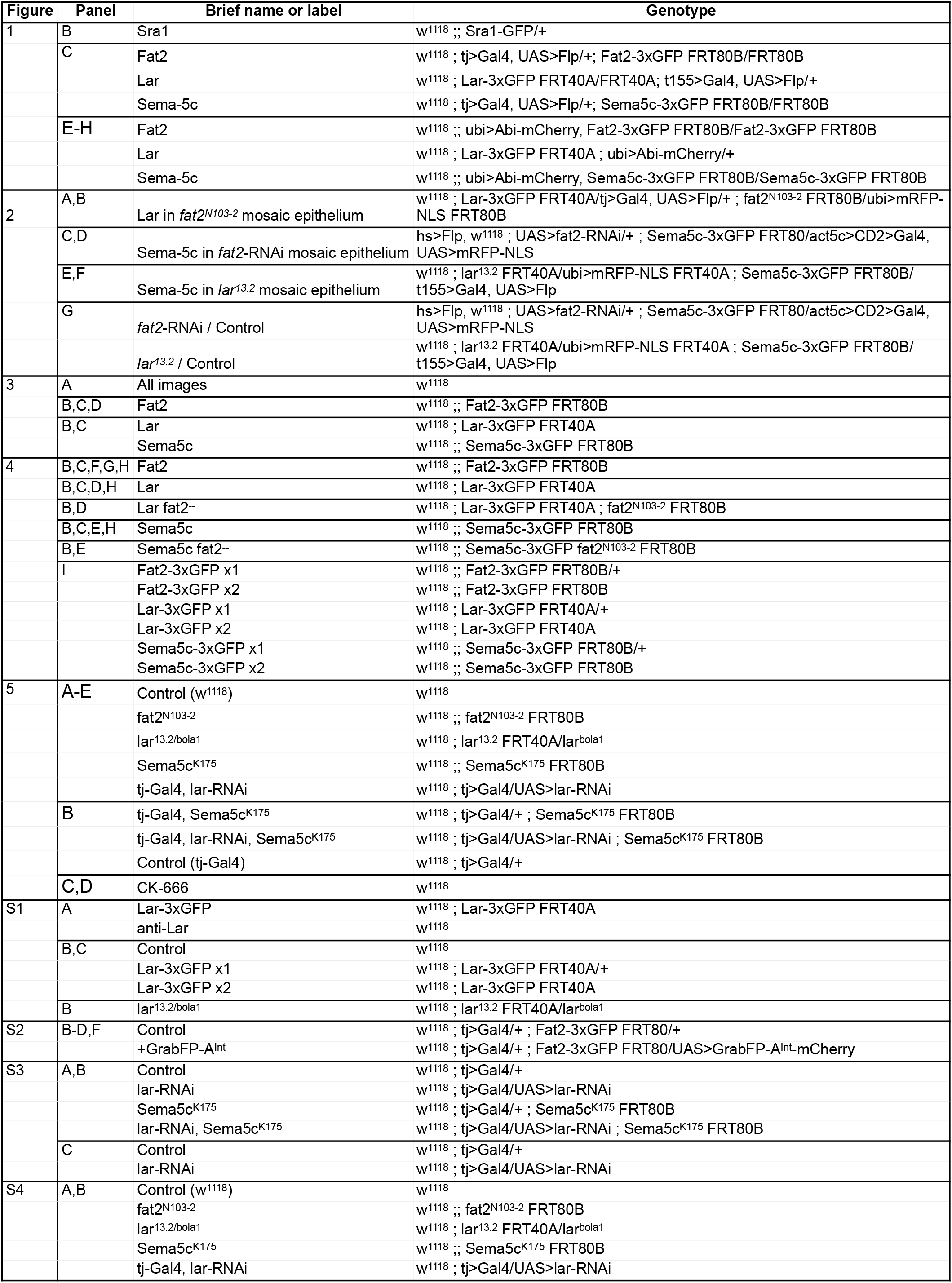
Genotypes of experimental animals

**Supp.Table 3:** *Drosophila* culture conditions for experiments

| Figure | Panel | Days on yeast | Temp (°C) |
| --- | --- | --- | --- |
| 1 | B | 2-3 | 25 |
|  | C - Fat2 | 3 | 25 |
|  | C - Lar | 5 | 25 |
|  | C - Sema5c | 3 | 25 |
|  | E-H | 2-3 | 25 |
| 2 | A,B | 3 | 25 |
|  | C,D | See methods for heat shock protocol |  |
|  | E,F | 5 | 25 |
|  | G | see C,D and E,F |  |
| 3 | A-D | 2-3 | 25 |
| 4 | B-G | 2 | 25 |
|  | H,I | 2-3 | 25 |
| 5 | A,C-E, B gray background (except tj-Gal4, lar-RNAi, Sema5c<sup>K175</sup> condition | 2 | 25 |
|  | tj-Gal4, lar-RNAi, Sema5c<sup>K175</sup> condition in A,C-E, B gray | 2 | 29 |
|  | B white background | 2 | 29 |
| S1 | A | 2-3 | 25 |
|  | B,C | 2 | 25 |
| S2 | B-D,F | 3 | 29 |
| S3 | A-C | 3 | 29 |
| S4 | A,C-E, B gray background (except tj-Gal4, lar-RNAi, Sema5c<sup>K175</sup> condition | 2 | 25 |
|  | tj-Gal4, lar-RNAi, Sema5c<sup>K175</sup> condition in A,C-E, B gray | 2 | 29 |

